# Creating cell-specific computational models of stem cell-derived cardiomyocytes using optical experiments

**DOI:** 10.1101/2024.01.07.574577

**Authors:** Janice Yang, Neil Daily, Taylor K. Pullinger, Tetsuro Wakatsuki, Eric A. Sobie

## Abstract

Human induced pluripotent stem cell-derived cardiomyocytes (iPSC-CMs) have gained traction as a powerful model in cardiac disease and therapeutics research, since iPSCs are self-renewing and can be derived from healthy and diseased patients without invasive surgery. However, current iPSC-CM differentiation methods produce cardiomyocytes with immature, fetal-like electrophysiological phenotypes, and the variety of maturation protocols in the literature results in phenotypic differences between labs. Heterogeneity of iPSC donor genetic backgrounds contributes to additional phenotypic variability. Several mathematical models of iPSC-CM electrophysiology have been developed to help understand the ionic underpinnings of, and to simulate, various cell responses, but these models individually do not capture the phenotypic variability observed in iPSC-CMs. Here, we tackle these limitations by developing a computational pipeline to calibrate cell preparation-specific iPSC-CM electrophysiological parameters.

We used the genetic algorithm (GA), a heuristic parameter calibration method, to tune ion channel parameters in a mathematical model of iPSC-CM physiology. To systematically optimize an experimental protocol that generates sufficient data for parameter calibration, we created simulated datasets by applying various protocols to a population of *in silico* cells with known conductance variations, and we fitted to those datasets. We found that calibrating models to voltage and calcium transient data under 3 varied experimental conditions, including electrical pacing combined with ion channel blockade and changing buffer ion concentrations, improved model parameter estimates and model predictions of unseen channel block responses. This observation held regardless of whether the fitted data were normalized, suggesting that normalized fluorescence recordings, which are more accessible and higher throughput than patch clamp recordings, could sufficiently inform conductance parameters. Therefore, this computational pipeline can be applied to different iPSC-CM preparations to determine cell line-specific ion channel properties and understand the mechanisms behind variability in perturbation responses.

**Author Summary:** Many drug treatments or environmental factors can trigger cardiac arrhythmias, which are dangerous and often unpredictable. Human cardiomyocytes derived from donor stem cells have proven to be a promising model for studying these events, but variability in donor genetic background and cell maturation methods, as well as overall immaturity of stem cell-derived cardiomyocytes relative to the adult heart, have hindered reproducibility and reliability of these studies. Mathematical models of these cells can aid in understanding the underlying electrophysiological contributors to this variability, but determining these models’ parameters for multiple cell preparations is challenging. In this study, we tackle these limitations by developing a computational method to simultaneously estimate multiple model parameters using data from imaging-based experiments, which can be easily scaled to rapidly characterize multiple cell lines. This method can generate many personalized models of individual cell preparations, improving drug response predictions and revealing specific differences in electrophysiological properties that contribute to variability in cardiac maturity and arrhythmia susceptibility.

**GLOSSARY:**

- **Model/parameter calibration**: tuning one or more parameters in the computational model so that the model output more closely matches experimental data
- **Experiment/protocol optimization**: the process of determining what type and amount of data is sufficient but also feasible for our model calibration goals
  - Protocol conditions – buffer calcium, potassium, or sodium concentrations; addition or removal of stimulus; pacing rates; channel block; etc.
  - Protocol length(?) – number of protocol conditions
  - Protocol data type(?) –AP, CaT, or both; normalized or non-normalized data
  - **Model prediction**: using the calibrated computational model to simulate response to *new (unseen)* conditions, drugs, or perturbations (in our case, I_Kr_ block)
  - Papers on independent validation/prediction

**Computational pipeline**: the full process of iPSC-CM computational model calibration; Includes iPSC-CM data acquisition/simulation -> data processing -> parameter calibration using genetic algorithm -> validation of calibrated models on an unseen condition (i.e. evaluating model predictions)

## INTRODUCTION

Human iPSC-derived cardiomyocytes (iPSC-CMs) are derived from patient cells that have been reprogrammed into pluripotency and subsequently manipulated to differentiate into the cardiac cell lineage. The development of this *in vitro* platform marked a technological breakthrough in cardiac research, and iPSC-CMs are now widely-used in cardiac pharmacology and disease research. iPSCs can be sourced through minimally-invasive procedures, such as skin biopsy or blood draw, and maintained or banked for long periods [1,2]. These properties of iPSC-CMs make them an ideal platform for pharmacological studies and personalized disease modeling [3]. However, electrophysiological variability between iPSC-CM preparations from different cell lines and differentiation methods limit the potential of this platform [4,5].

Mathematical models of cardiomyocyte electrophysiology can provide valuable insight into iPSC-CM electrophysiological variability. These models contain parameters describing the ionic currents and fluxes that contribute to the properties of cardiac action potential (AP) and calcium transient (CaT) waveforms. Several models of iPSC-CM physiology exist in the literature [6–8], with the most recent and comprehensive model published by Kernik et al. in 2019, hereon referred to as the “Kernik model” [9]. However, the baseline parameter values in these models are not representative of the electrophysiological heterogeneity observed in iPSC-CMs. Additionally, parameter calibration has often relied on separate patch clamp measurements of each type of current. This process is time-consuming, inaccessible to many research groups, and often captures the physiology of only the average behavior of the population of cells examined. Methods to simultaneously optimize several or all model parameters using minimal data are under development [10–14]. Recent work on guinea pig ventricular myocyte models [13] and human iPSC-CM models [14] have demonstrated the utility of automated tuning algorithms for improving model parameterization. For example, in Devenyi et al [13], an adjusted model, constrained by data, predicted the effect of slow delayed rectifier K^+^ current (I_Ks_) block more accurately compared to baseline model. Analogous work, at the level of individual ion channels, has shown the superiority of sinusoidal voltage clamp protocols, compared with traditional square-wave protocols, for calibration of channel kinetic parameters [15].

Even with recent advances in iPSC-CM maturation protocols [16–19], most iPSC-CMs still display embryonic- or neonatal-like cardiac phenotypes with different AP and CaT shapes compared with adult myocytes. These phenotypic differences, and how they change temporally, can depend on subtle differences in cell culture conditions and the genetic background of the cell donor [4,5,20], which likely contributes to the wide variation observed between studies [4,5]. No standardized procedure currently exists for characterizing iPSC-CM heterogeneity; prior efforts have generally examined a handful of molecular markers [21], used subjective observations of physiology [22,23], or employed specialized methods such as single cell RNA sequencing [24,25]. These factors, and the potential importance of iPSC-CMs as a research platform, highlight the need for automated methods to characterize the cell lines used in each study.

Here we combined these ideas of simultaneous calibration of multiple parameters and optimization of a minimal experimental protocol for generating calibration data. We aimed to create digital twins of iPSC-CM cell preparations from the Kernik iPSC-CM model, thereby connecting observed iPSC-CM electrophysiology with molecular function and mechanisms. We hypothesize that fluorescence readouts of iPSC-CM physiology under varied experimental conditions provide enough information to reveal cellular ion channel properties, which can then be incorporated into the iPSC-CM digital twins. To systematically evaluate various data types and experimental protocols in their ability to inform model parameters, we created a simulated, *in silico* dataset from Kernik model variations with known parameter values. From this study, we developed a computational pipeline which includes: 1) an optimized protocol for fluorescence recordings of iPSC-CM preparations; 2) a genetic algorithm process for calibration of ion channel parameters in iPSC-CM computational model, using the experimental recordings; and 3) validation of the resulting calibrated models by evaluating their predictions on independent, yet physiologically-important, perturbations. We demonstrate the utility of our computational pipeline in generating iPSC-CM digital twins that capture ionic variability and predict cell-specific electrophysiological phenotypes, including drug-induced arrhythmia susceptibility.

## RESULTS

Our primary objective was to optimize an experimental protocol for generating sufficient data to calibrate an iPSC-CM mathematical model, taking experimental feasibility and throughput into account. To systematically evaluate how different types of data and experimental conditions impact parameter calibration and model predictions of responses to new conditions or drug treatments, we created a simulated *in silico* dataset, generated from a population of Kernik models with random variations in their 16 maximal conductance parameters (Figure 1A, Supplementary Table S2). We simulated AP and CaT generated by these model cells under various conditions (Supplementary Table S3), and then concatenated these data in various combinations based on the “candidate protocols” we wanted to evaluate. The 16 Kernik model conductance parameters were then calibrated, using a genetic algorithm, to best match the steady state AP and CaT traces from each candidate protocol. These fitted parameters were incorporated into the Kernik model, and the newly-calibrated models were evaluated on their ability to predict the original model cell’s responses to new conditions (outlined in <u>methods</u> and Figure 1B). Optimizing the experimental protocol using this *in silico* dataset provided 3 major advantages: 1) knowledge of the ground truth parameter values that generated the dataset, so the accuracy of the calibrated parameters can be evaluated, 2) relatively fast and easy simulation of new conditions from the same or additional model cells, in comparison to acquiring new in vitro data and cell lines, and 3) generating corresponding data types through computational methods and data processing, forgoing the need to set up new experiments to collect each data type.

**Figure 1:**
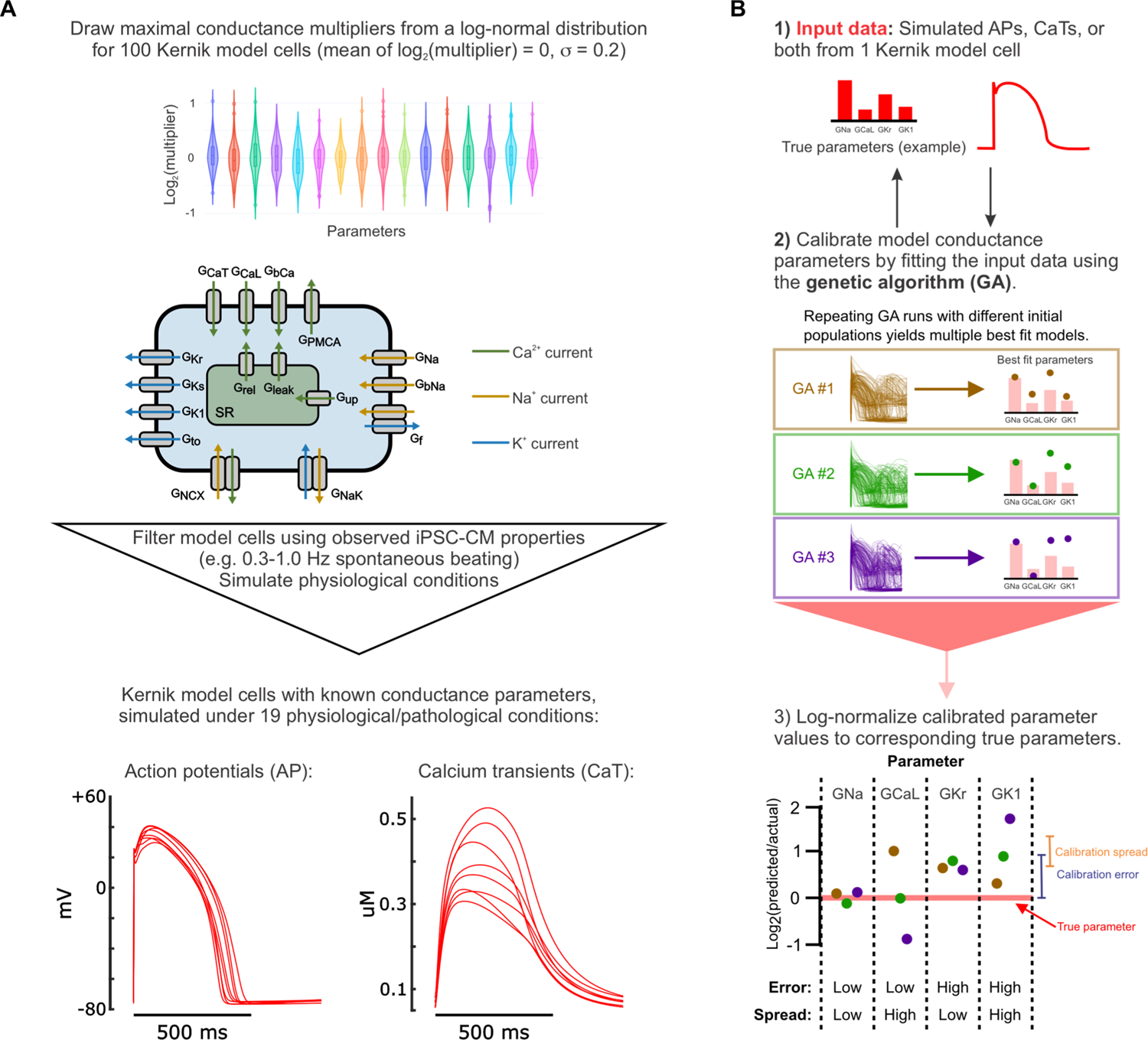
Generation of *in silico* dataset and calibration pipeline optimization. (A) Schematic of the workflow for creating the *in silico* dataset used to optimize the calibration pipeline. (B) Schematic of the genetic algorithm (GA) parameter calibration process, including the true parameters that generated the input data (1), multiple genetic algorithm runs to fit this data (2), and metrics used to evaluate the resulting fitted parameter values (3). The genetic algorithm (GA) creates a population of model cells and simulates the same conditions that generated the input data. Then, the error between GA-generated traces and input data is calculated to determine how to modify the population, creating a new population. This process is repeated until the population’s errors converge (∼20 iterations). For parameter evaluation: calibration error = | log_2_(fitted parameter value / true parameter value) |; calibration spread = standard deviation of log_2_(fitted parameter value / true parameter value) between 10 GA runs.

### Simultaneously calibrating to AP and CaT data improves model predictions

First, we examined whether AP or CaT recordings alone provided sufficient information for conductance parameter calibration and model predictions. We used simulated recordings from baseline physiological conditions without pacing stimuli (spontaneous APs), and supplied to a genetic algorithm (GA) either 1) only the AP traces, 2) only the CaT traces, or 3) both (Figure 2A). The GA calibrates parameters by varying them to create a population of Kernik model cells, then iteratively selecting and modifying individual models that generated AP and CaT traces that best matched the supplied data (outlined in Figure 1B) [26]. We ran the genetic algorithm 10 times per input dataset, with each GA run starting on a different initial population of models. This allowed us to assess whether each input dataset provided enough information to consistently estimate parameter values, even when the initial search space varied.

**Figure 2:**
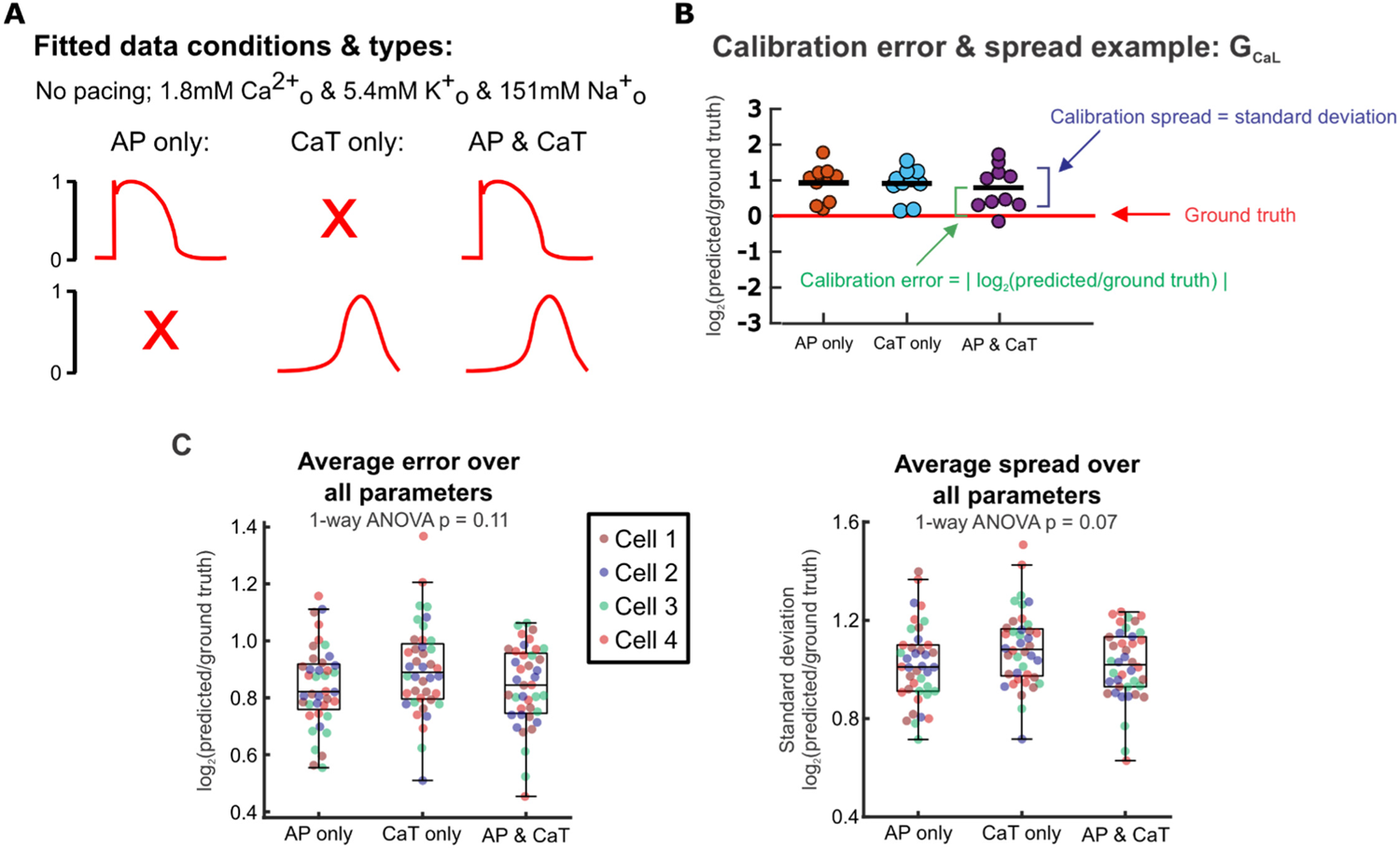
Calibration error and spread remain similar whether only AP, only Ca transient, or both are included during calibration. (A) Schematic of fitted data types. (B) Example of calibrated parameter values from fitting each data type for one parameter (G_CaL_), log-normalized to the ground truth G_Cal_ value of the fitted model cell. Parameter error and spread calculations are visualized. (C) Parameter errors and spreads calculated by comparing calibrated parameters to the corresponding true parameters of 4 model cells from the *in silico* dataset. Each point represents the average error or spread over all 16 fitted parameters from a single genetic algorithm run for 1 model cell.

We evaluated each candidate protocol on parameter calibration accuracy (calibration error) and consistency (calibration spread). Calibration error was represented by the absolute values of calibrated value log-normalized to the corresponding ground truth parameter value, and calibration spread was assessed by calculating the standard deviation (spread) in log-normalized calibrated parameter values between the 10 runs (Figure 2B). After averaging over all 16 fitted parameters, we found minimal differences in calibration error (p = 0.11) or spread (p = 0.07) between fitting only voltage, only calcium, or both simultaneously (Figure 2C). From these initial tests, therefore, there appears to be little difference between experimental protocols in how well (or poorly) each can identify model parameters.

We next assessed the calibrated models from each candidate protocol on how well each predicted model cell responses to 30% block of rapid delayed rectifier current I_Kr_,as this perturbation is both: 1) independent of the conditions used for model calibration and 2) relevant to cardiac pharmacology [27]. With this evaluation (Figure 3A), we found that models fitted to only AP data or only CaT data differed significantly from the ground truth model (Figure 3B, Supplementary Figure S1). When both AP and CaT recordings were used simultaneously during parameter calibration, the resulting calibrated models exhibited substantial improvement in their predictions of 30% I_Kr_ block. Therefore, fitting parameters to both AP and CaT traces simultaneously is required for accurate prediction of results that were not used in the fitting process.

**Figure 3:**
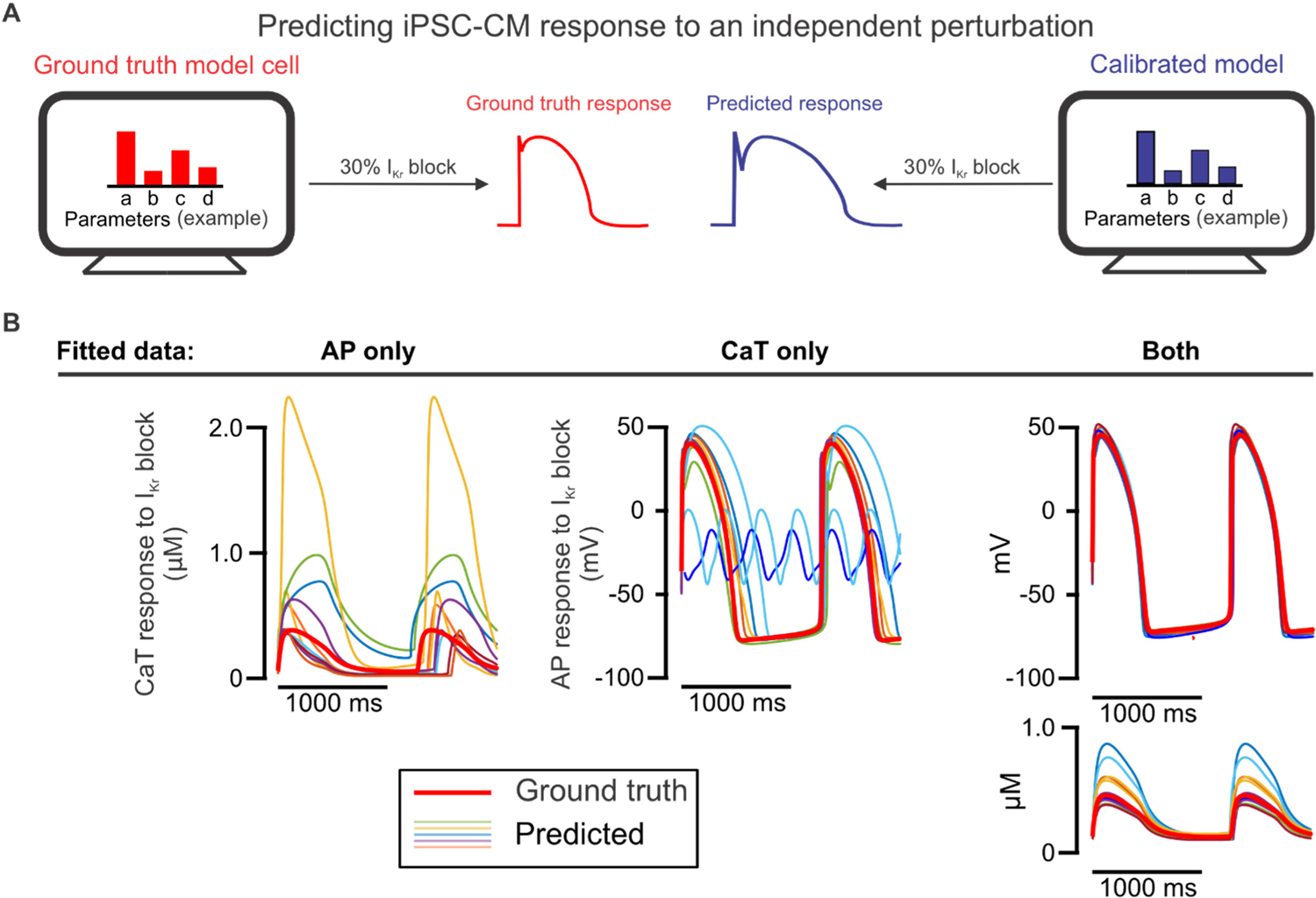
Prediction of iPSC-CM response to an independent perturbation differentiates the models calibrated to AP and CaT simultaneously from those calibrated to single traces. (A) Schematic illustrating the evaluation of calibrated models on their ability to predict an independent response. (B) AP and/or CaT responses to 30% l_Kf_ block predicted by 10 calibrated models for one model cell (thinner traces), compared with the true response of the corresponding model cell (thicker red trace).

### Overall parameter error and spread show minimal change with varied protocol conditions

Next, we explored how varying the number and identity of simulated conditions in the candidate protocols affected parameter calibration. AP and CaT recordings were simulated under many conditions that could be produced in an electrophysiology laboratory and constructed candidate protocols from various combinations of these conditions, which included: 1) varied buffer calcium, from hypo- to normal to hyper-calcemic conditions, 2) spontaneous beating and varied pacing rates from 1-2 Hz, 3) varied levels of L-type calcium channel (I_CaL_) blockade, and 4) a mixture of these conditions (see Methods for details). We also separated these data into shorter protocols, to see if the experimental setup could be further simplified (Figure 4A). Unexpectedly, we found that parameter errors and spread across all 16 maximal conductance parameters did not differ significantly between the protocols with varied cell culture conditions (Figure 4B, p = 0.814 and p = 0.673, respectively). Additionally, we observed no significant differences in average calibration errors (p = 0.088) and spreads (p = 0.137) when varying the number of conditions in the protocol (Figure 4C). Similar findings were observed when evaluating different combinations of these experimental conditions (Supplementary Figure S2).

**Figure 4:**
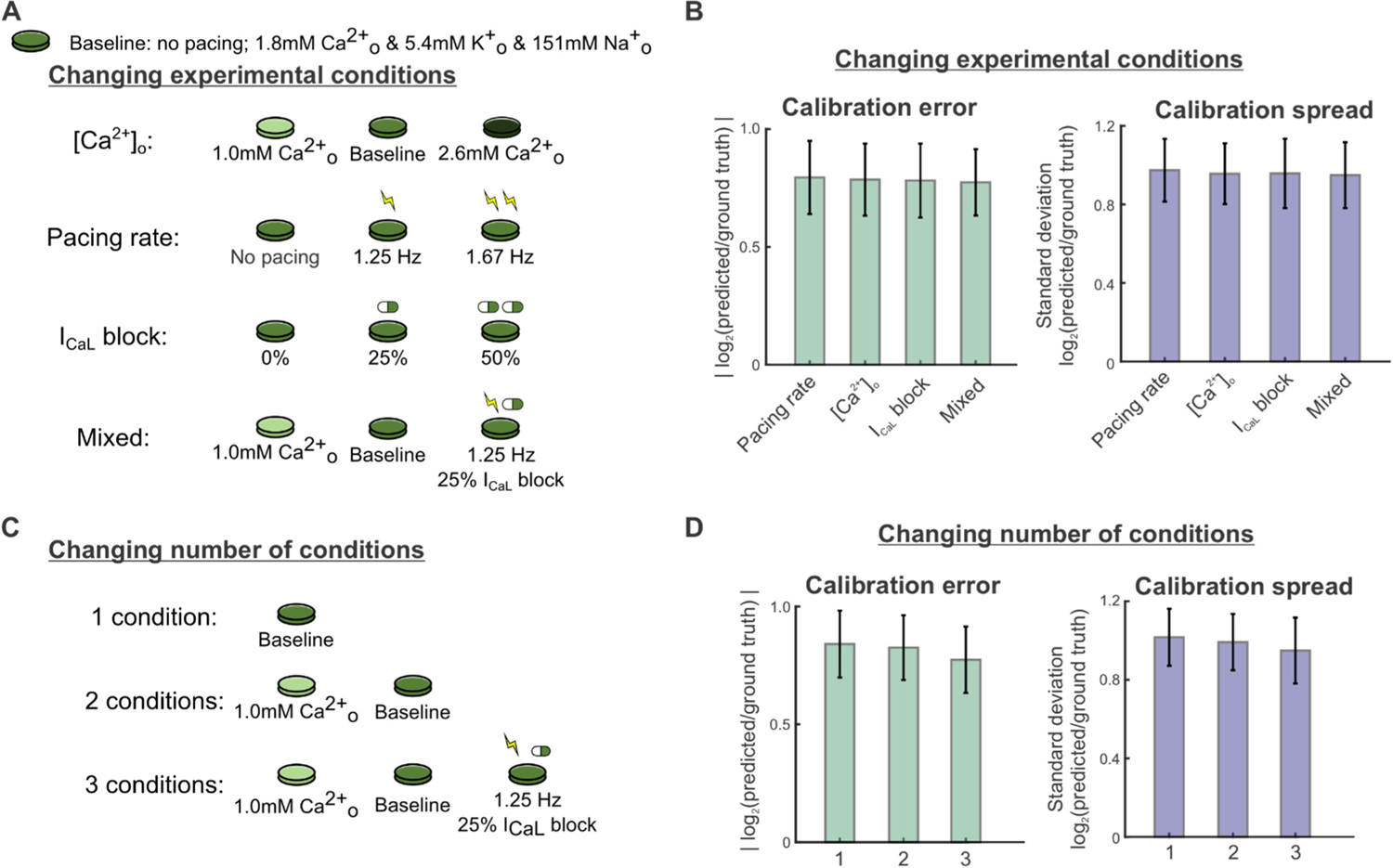
Varying the types and number of experimental conditions has minimal effect on overall calibration error and spread. (A) Schematic of candidate protocols to generate data for model calibration: varied buffer calcium concentration, varied pacing rates, varied l_CaL_ block levels, and a mixture of the 3. (B) Parameter errors and spreads from calibration runs for 4 Kernik model cells, using calibration data from each protocol in Fig. 4A. Bar plots show the mean calibration errors and spreads over all 16 fitted parameters from 40 calibration runs (10 runs per model cell). Error bars represent standard deviations over these runs. (C) Schematic of candidate protocols to generate data for model calibration: combinations of 1, 2, or 3 parts of the mixed condition protocol. (D) Parameter errors and spreads from calibration runs for 4 Kernik model cells, using calibration data from each protocol in Fig. 4B.

### A readily-achievable 3-condition protocol improves predictions of I_Kr_ block response

As with the results shown in Figure 3, we assessed how well calibrated models predicted the response to I_Kr_ block, and we found that accuracy depended strongly on the experimental conditions simulated in each candidate protocol (sample AP traces shown in Figure 5A). Out of the protocols using 3 experimental conditions in Figure 4A, the mixed-conditions protocol generated calibrated models with the most accurate predictions of changes in APs with 30% I_Kr_ block, compared with protocols which varied only the buffer calcium, pacing rates, or I_CaL_ blockade (Figure 5B-C). When evaluating whether a shorter version of this protocol could produce equally predictive models, the full protocol with all 3 conditions outperformed protocols with 1 or 2 of the conditions (Figure 5B-C). These data suggest that an optimal experimental protocol for iPSC-CM model parameter calibration should include voltage and calcium transient fluorescence recordings under at least 3 varieties of cell culture condition changes or perturbations. Of note, this supports our hypothesis that data from iPSC-CMs under a variety of conditions provides sufficient parameter identification to generate predictive models for pharmacological applications.

**Figure 5:**
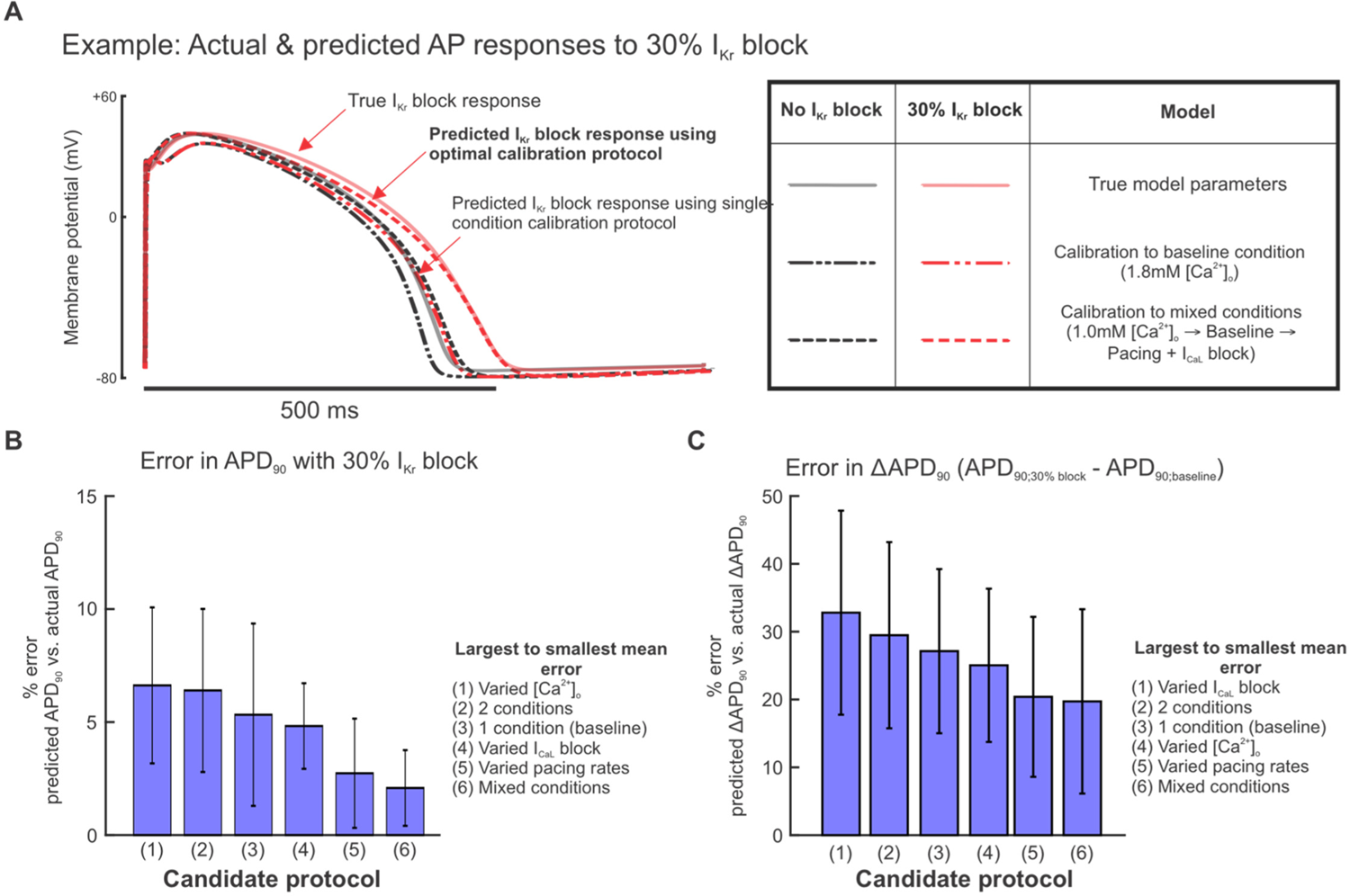
Calibrating parameters to data from the mixed condition protocol improves model predictions of l_Kr_ block response. (A) Examples of simulated baseline (black graces) and 30% l_Kr_ block (red traces) steady state APs using the ground truth model cell (solid traces) and predicted models (dashed traces) from calibrations to the indicated protocols. (B) Percent error in calibrated model predictions of APD_90_ values at 30% l_Kr_ block. Bars represent means over 10 calibration runs and error bars represent standard deviations. APD_90_ values were normalized to corresponding AP amplitudes before calculating percent error. (C) Percent error in calibrated model predictions of change in APD_90_ change between baseline and 30% l_Kr_ block. APD_90_ values were normalized to corresponding AP amplitudes before calculating percent error.

### Normalized fluorescence recordings sufficiently inform parameter calibration

Fluorescence recordings generally detect relative changes and do not provide absolute levels of voltage and calcium. Fluorescent dyes can be calibrated [28,29], and patch clamp can detect true transmembrane potentials, but these are more challenging and lower-throughput techniques, compared with straightforward fluorescence recordings. To assess the potential implications for parameter identification, we compared two versions of simulated recordings obtained with our optimized protocol: 1) unscaled (original) *in silico* data, representing patch clamp and calibrated calcium measurements, and 2) normalized versions of the same waveforms, representing simpler fluorescent recordings (Figure 6A).

**Figure 6:**
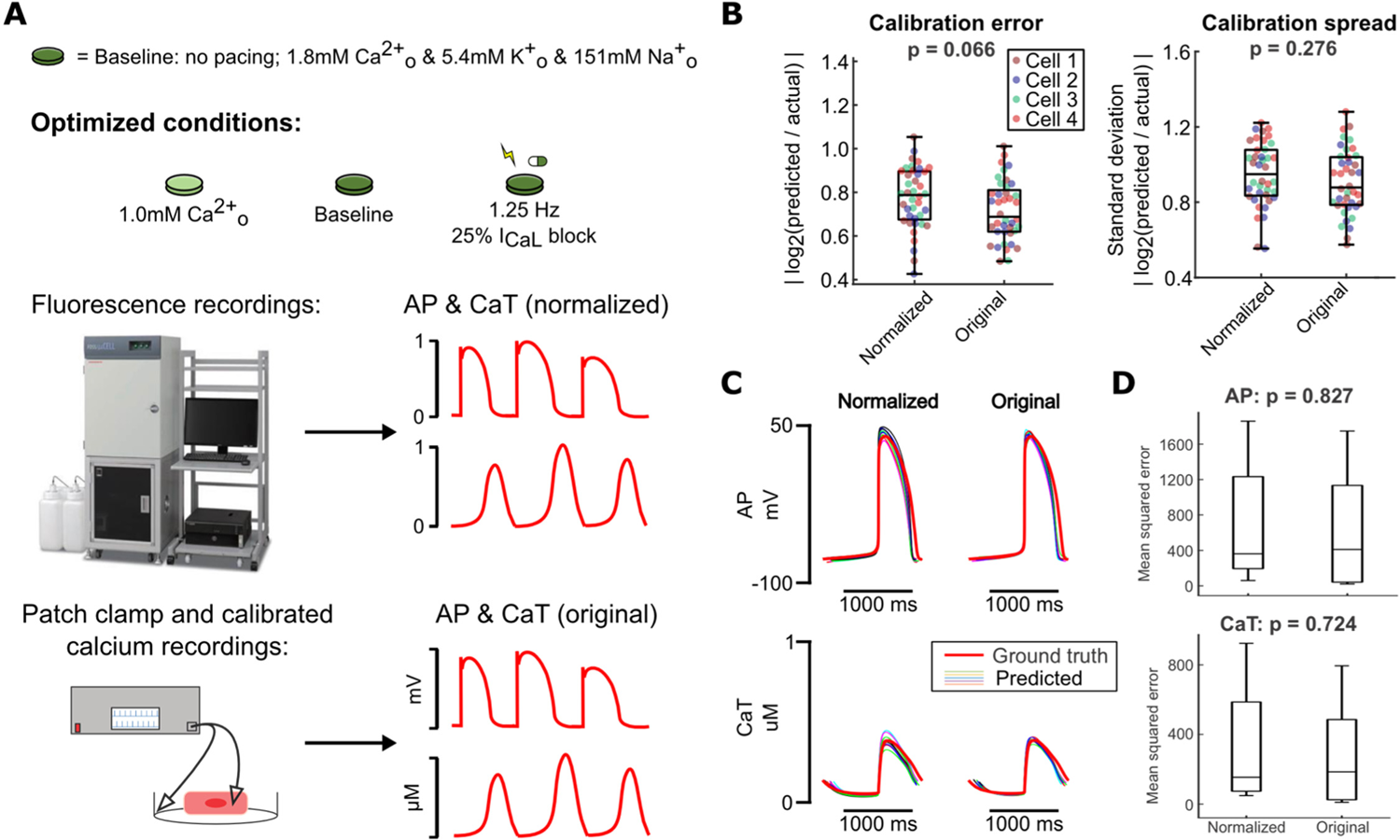
Normalization of calibration data does not significantly affect model calibration results. (A) Schematic of optimized experimental conditions, normalization of simulated data used to represent fluorescence recordings, and original simulated data used to represent more precise measurement modalities. (B) Parameter errors and spreads from calibration runs on data from 4 Kernik model cells, using either normalized or original data for calibration. Each point represents the average error or spread over all fitted parameters. (C) Predicted AP and CaT responses to 30% l_Kr_ block (thinner traces) from each set of conditions, compared to the true response of the corresponding model cell (thicker red trace). (D) Mean squared errors between data points in the predicted traces (simulated using calibrated models) and the actual model cell’s traces, for simulations under 30% l_Kr_ block.

Parameter calibration accuracy and consistency did not differ significantly when comparing calibrated models from the original data with those from the corresponding normalized data (Figure 6B). Notably, calibrated models from these two data types also performed similarly when evaluating predictions of 30% I_Kr_ block response (Figure 6C). This suggests that fluorescence voltage and calcium recordings from our optimized protocol conditions provide sufficient information to calibrate predictive models.

### Calibrated models predict variability in arrhythmia susceptibility *in silico*

The primary goal of optimizing a computational pipeline for model calibration is to create cell preparation-specific models that can predict variability in phenotypes, particularly arrhythmia susceptibility, between iPSC-CM lines. To assess the ability of our optimized protocol to predict cell line-specific arrhythmia susceptibility, we calculated the lowest level of I_Kr_ block (i.e. highest % I_Kr_) that induced arrhythmia dynamics (afterdepolarizations, Torsades de Pointes, alternans, beating cessation, or tachycardia) in the same 4 Kernik model cells from our *in silico* dataset. These simulations defined, for each of the 4 model cells, an “I_Kr_ block tolerance threshold” (Figure 7A), which ranged from 37% I_Kr_ block (highest susceptibility) to 62% I_Kr_ block (highest tolerance) (Figure 7C) across the 4 model cells.

**Figure 7:**
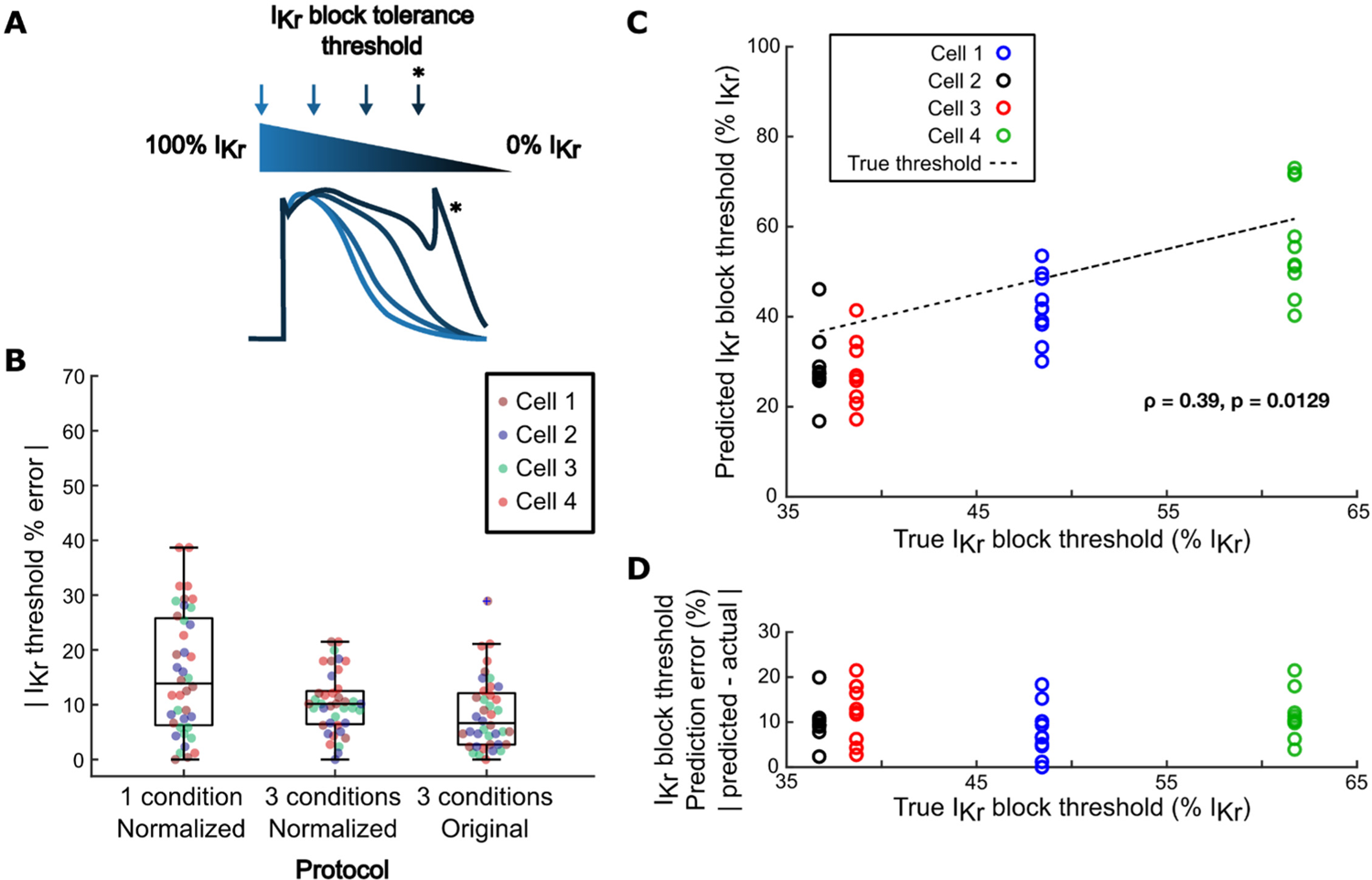
Models generated by the optimized calibration pipeline predict cell-specific arrhythmia susceptibility. (A) Schematic of calculation of l_Kr_ block tolerance threshold. (B) Errors in predictions of l_Kr_ block tolerance threshold, comparing calibrated models from 3 protocols: 1) normalized AP and CaT recordings from a single physiological condition; 2) normalized AP and CaT recordings from the optimized, 3-condition protocol; and 3) non-normalized AP and CaT recordings from the optimized protocol. (C) Correlation between true l_Kr_ block threshold of model cells and their corresponding calibrated models’ predicted l_Kr_ block thresholds. (D) Correlation between true l_Kr_ block threshold of model cells and their corresponding calibrated models’ l_Kr_ block threshold prediction error.

Predicted I_Kr_ block tolerance thresholds were also calculated for the calibrated models generated by our computational pipeline. These predicted thresholds were assessed against the ground truth threshold of the model cell. Calibrated models from our computational pipeline predicted I_Kr_ block thresholds with high accuracy and consistency, with similar error and spread as models calibrated to corresponding non-normalized recordings (Figure 7B). The aggregated calibrated models also predicted relative arrhythmia susceptibility of the 4 model cells, as shown by a significant positive correlation between ground truth and predicted I_Kr_ block thresholds in Figure 7C. Additionally, there was no correlation between the model cell’s true I_Kr_ block threshold and its associated calibrated models’ prediction error, suggesting that the accuracy of these predictions would not be affected by variability in arrhythmia susceptibility between iPSC-CM preparations (Figure 7D).

### Optimized model calibration process constrains key ionic parameters

Throughout the protocol optimization process, we observed that while calibration error and spread largely remained constant when averaged over all fitted parameters, certain individual parameters consistently showed low calibration spread (e.g. G_Kr_, G_NaK_) or high calibration spread (e.g. G_rel_, G_CaT_) (Supplementary Figure S3). Thus, we hypothesized that our calibration method only needs to tightly constrain a few key parameters, tolerating inaccuracy or variation in less important parameters while still generating highly predictive calibrated models. To determine whether normalized data from the optimized protocol constrains the parameters that play the largest roles in determining AP and CaT morphology, we analyzed the relationship between parameter calibration accuracy and consistency with the Kernik model’s sensitivity to parameter changes. We used multivariable regression to determine how the 16 calibrated model parameters affect relevant phenotypes such as AP duration, CaT amplitude, and I_Kr_ block threshold (Figure 8A) [30]. The resulting model coefficients represent the magnitude and direction of the effect of changes in the corresponding parameter on the modeled phenotype (Figure 8A-C). The most important parameters depended on which model output was considered (Figure 8A-C), but parameters that appeared frequently included rapid delayed rectifier K^+^ conductance (G_Kr_), L-type Ca^2+^ conductance (G_CaL_), Na^+^ conductance (G_Na_), inward rectifier K^+^ conductance (G_K1_), and Na^+^/K^+^ pump conductance (G_NaK_).

**Figure 8:**
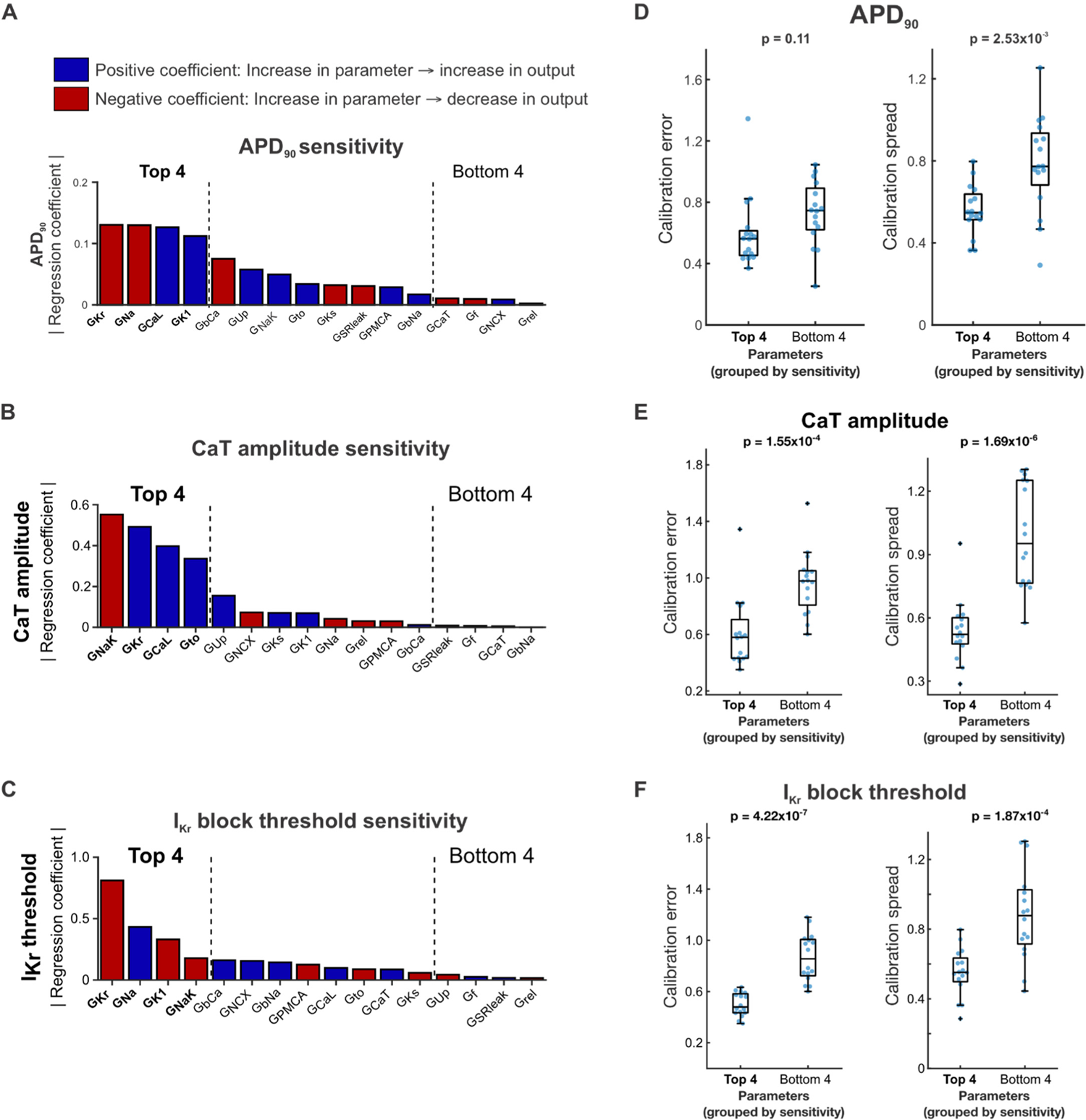
Highly sensitive conductance parameters are well-constrained by the optimized calibration pipeline. (A-C) Regression coefficient magnitudes for each Kernik model maximal conductance parameter’s contribution to (A) APD90, (B) CaT amplitude, and (C) l_Kr_ block threshold. Blue bars indicate positive coefficients, and red bars indicate negative coefficients. The top 4 and bottom 4 parameters, ranked by regression coefficient magnitude, are bolded and indicated with black dashed lines. (D-F) Distributions of parameter calibration errors (left) and calibration spread (right) from the optimized calibration pipeline, with parameters grouped into the top 4 and bottom 4 regression coefficient magnitudes for (D) APD90, (E) CaT amplitude, and (F) l_Kr_ block threshold. Each blue dot represents the calibration error or spread value for 1 parameter from calibrations on one model cell (4 parameters per group x 4 model cells =16 points).

We then grouped the parameters into the highest and lowest regression coefficient magnitudes for each phenotype, and assessed calibration errors and calibration spreads from the optimized calibration protocol within those groups. The 4 parameters with highest regression coefficient magnitudes for CaT amplitude and I_Kr_ block threshold showed significantly lower average calibration error and spread compared with the 4 parameters with the lowest regression coefficient magnitudes (Figure 8E-F). Accordingly, the optimized calibration protocol displayed significant improvement over most other candidate protocols in predictions of CaT amplitude and I_Kr_ block threshold. We saw similar results when examining calibration errors and spreads in parameters grouped by APD_90_ sensitivity, though the difference in calibration errors was not statistically significant (Figure 8D). These results suggest that our optimized computational pipeline does constrain the key Kernik model conductance parameters needed to generate predictive models.

### Validation of optimized computational pipeline on *in vitro* iPSC-CM recordings

After optimization of our computational pipeline using *in silico* data, where the ground truth model parameter values were known, we assessed whether our pipeline could still generate predictive models using *in vitro* data. We applied the computational pipeline to normalized fluorescence AP and CaT recordings from one iPSC-CM cell line under different combinations of 3 varied conditions: 1) low buffer calcium (1.0 mM) with 1 Hz pacing, 2) physiological buffer calcium (1.8 mM) with 1 Hz pacing, and 3) physiological buffer calcium with 1.25 Hz pacing. Figure 9A outlines the data processing and normalization procedures performed on the fluorescence recordings prior to model calibration. Recordings from a separate condition (1.0 mM buffer calcium with 2 Hz pacing) were left out from model calibration, so they could be used to independently evaluate predictions generated by the calibrated model. Calibrating the model conductances using data from only one of these conditions resulted in unconstrained conductance values and highly variable predictions of response to the left-out validation data (Figure 9B).

**Figure 9:**
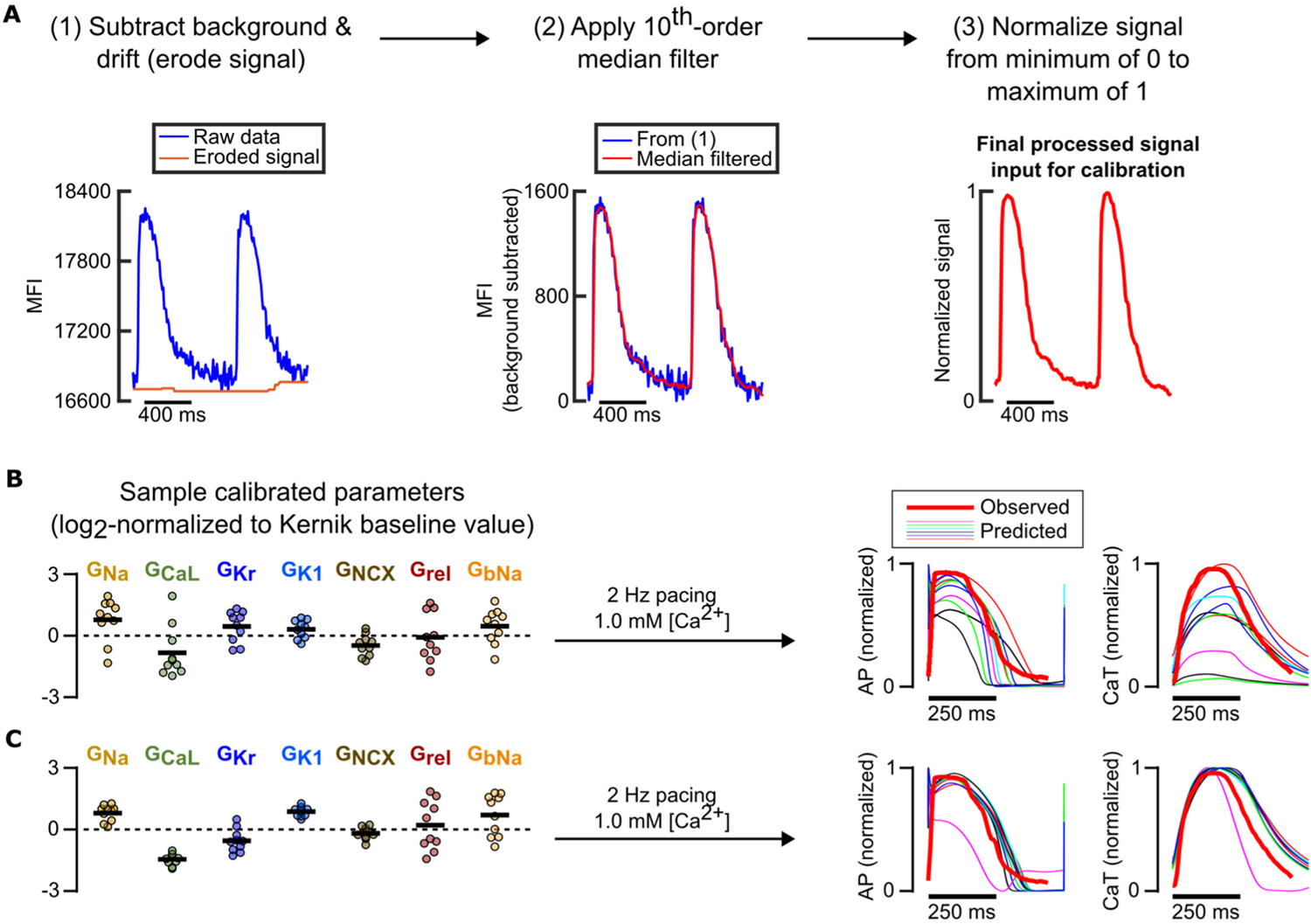
Preliminary model calibrations to *in vitro* data support findings from *in silico* calibrations. (A) Fluorescence recording processing steps in preparation for model calibration pipeline. (B) Resulting calibrated conductance values (left) normalized to Kernik model baseline values (dashed line at 0) and predicted traces under an independent condition (right) when only data from a single condition (physiological buffer, 1.25 Hz pacing) is used to fit parameters. (C) Resulting calibrated conductance values (left) normalized to Kernik model baseline values (dashed line at 0) and predicted traces under an independent condition (right) when data from varied pacing rates and buffer [Ca^2^*] are used to fit parameters.

Including data from 2 additional, mixed conditions significantly improved consistency of key conductance parameter calibrations (Figure 9C). When these calibrated parameters were incorporated into the Kernik model and the cell line’s response to a new condition was predicted, we found that models calibrated to recordings from 3 conditions substantially outperformed models calibrated to only 1 condition. These results further supported our findings from *in silico* protocol optimization, showing that data from fluorescence voltage and calcium transients under 3 varied conditions can constrain key model parameters and generate predictive, cell preparation-specific models.

## DISCUSSION

While the application of iPSC-derived cardiomyocytes to cardiac disease and pharmacology research has grown dramatically in the past decade, intrinsic and extrinsic variability between cell preparations still hinders reproducibility and clinical translation of studies that use these cells. Generic out-of-box mathematical models of iPSC-CMs do not reflect this variability and potentially generate inaccurate predictions when used to simulate pharmacological ion channel effects or genetic variations [10]. In this study, we aimed to overcome these limitations of the iPSC-CM *in vitro* platform and their mathematical models by creating a computational pipeline which can rapidly identify the cell preparation-specific ionic properties contributing to this phenotypic variability. The parameter calibration algorithm we used, a heuristic genetic algorithm, allows for selection of specific parameters of interest or parameter value limits, and is suitable for parallel computing. To determine the feasibility of this approach, we used a model-generated *in silico* dataset to evaluate candidate protocols containing varied data types and experimental conditions. We found an optimized protocol for fluorescence AP and CaT recordings, consisting of varied buffer calcium concentration, electrical pacing, and I_CaL_ block conditions, which provided enough information to calibrate accurate and predictive cell-specific models, while remaining short in duration and straightforward to acquire. Subsequent model calibrations with an *in vitro* dataset suggested potential flexibility in which specific conditions are selected for the protocol. In our computational pipeline development and subsequent validation, these data were able to inform key ionic contributors, generating calibrated models which accurately and consistently predicted cell-specific responses to I_Kr_ block.

### Calibration of models of cardiac electrophysiology

Parameter values in computational models of cardiac electrophysiology are often determined by manual fits to voltage and current recordings, often collected under one or few experimental conditions. This limits the baseline models’ ability to represent electrophysiological heterogeneity. Additionally, determination of these baseline parameter values may be affected by selection bias for cells with larger currents, inadequate separation of activation and inactivation kinetics, and inter-laboratory differences in current and voltage clamp protocols [31]. These issues have, in many cases, led researchers to recalibrate the model parameters in attempts to better reflect the behaviors of their specific cardiomyocyte preparations. For example, Potse et al. fitted a reaction-diffusion model of ventricular electrical activity to reproduce phenotypes of patients with heart failure or left branch bundle block [32]. Similarly, Lombardo et al. tuned atrial electrophysiology models for patients undergoing ablation therapy [33] whereas Krogh-Madsen et al. used the genetic algorithm to fit a human ventricular cardiac model to clinical QT interval data [34]. In all three cases, these groups found significant discrepancies between the respective initial models and their final, calibrated, fit-for-purpose models. However, to the best of our knowledge, the experimental data used in virtually all previous cardiomyocyte model calibration research come from voltage clamp, patch clamp, or clinical data. Currently, these data are still difficult to obtain from a large group of patients or cell preparations. More recent work has demonstrated the utility of genetic algorithms and the resulting recalibrated models in prediction of arrhythmic behaviors and drug mechanisms, but these also used complex voltage step protocols to calibrate model parameters [35]. These prior results motivated our search for protocols that could calibrate mathematical models using voltage- and calcium-sensitive fluorescent dye measurements, which are considerably easier to acquire than patch-clamp recordings.

### Guiding experimental design for optimal parameter calibration

We considered several factors when optimizing the experimental protocol to generate iPSC-CM data for our computational pipeline: 1) the information that the data would provide for parameter calibration, 2) the complexity of the protocol, and 3) the feasibility of the experiment for broad accessibility and high throughput setups. Since the goal of model calibration is usually to improve the accuracy of model parameterizations and predictions, previous work has largely focused on designing experiments to maximize the information content of the resulting data [36–38]. In particular, for cardiac electrophysiology models, model calibration has almost universally been performed on data from complex voltage step protocols, in order to accurately determine individual ion channel conductances and kinetics [10,39,40]. Here, we emphasize that it is also important to identify adaptable protocols that can be widely adopted. Therefore, we opted to focus on using fluorescence recordings to calibrate model parameters, instead of the usual techniques such as patch clamp. To evaluate different protocols, we created an *in silico* dataset for testing various data combinations in their ability to inform parameter calibration. This approach provided 2 major advantages. First, guided by experimental feasibility and relevance, we were able to simulate iPSC-CM electrophysiology under many combinations of conditions without the time and financial costs associated with *in vitro* experiments. Second, we knew the ground truth ionic parameter values that produced these AP and CaT traces, so we could precisely assess which simulated conditions informed which parameters. We also ran multiple rounds of genetic algorithms on each tested *in silico* protocol, with each round starting from a different initial model population, to assess how consistently each parameter was identified. We used this as a measure of confidence in each parameter estimate.

In their analysis on model parameter constraint using regression and Bayesian approaches, Sarkar and Sobie found that, in both cases, parameter constraint improved when values for more than one output feature were provided [41]. Similarly, we found that calibrating model parameters to fluorescence recordings of AP and CaT traces, as opposed to using only one of the two, resulted in better parameter constraint for some (but not all) parameters. Perhaps more importantly, we found that calibrating to both outputs simultaneously resulted in significant improvement of the calibrated models’ predictions on a new, independent output. This is consistent with prior results which showed that, when translating drug responses across cell types, using multiple electrophysiological features or observations of iPSC-CMs under multiple conditions improved translations [42–44]. Similarly, our final optimized protocol included a mixture of 3 experimental conditions. Finally, overall parameter calibration accuracy and consistency, as well as independent predictions of I_Kr_ block response, were similar between the models fitted to normalized data (representing fluorescence data) and those fitted to corresponding non-normalized data (representing patch clamp and calibrated Ca^2+^ transient data). Collectively, these findings support our hypothesis that data from fluorescence recordings under multiple conditions, which are more practical compared with microelectrode or patch clamp recordings, can be used to calibrate predictive models. Notably, several parameters such as the background Ca^2+^ and Na^+^ conductances, G_Ks_, and G_rel_ were rarely constrained by any of the datasets we tested. Our subsequent parameter sensitivity analysis showed that model conductances that strongly affect relevant model outputs tend to be well-constrained by our optimized protocol.

### Prediction of cell-specific arrhythmia susceptibility

Finally, we demonstrated that our optimized computational pipeline can generate consistent and predictive mathematical models from *in vitro* iPSC-CM data, using fluorescence voltage and calcium recordings under a similar set of experimental conditions. Many therapeutics have the potential to induce patient-specific arrhythmic effects through their intended targets (e.g. quinidine blockade of I_Na_ and I_Kr_) or unintended effects (e.g. antibiotic off-target blockade of K_+_ channels) [45,46]. Previous analyses of systems biology models have also argued the importance of a prediction-focused computational modeling approach. In an influential paper, Gutenkunst et al. posited that a model that accurately predicts relevant phenotypes even with some parameter value inaccuracies is more useful than a model with high parameter accuracy and consistency, but inaccurate predictions [47]. This suggests that attempting to determine parameter values directly through precise measurements is an inefficient way of optimizing models. Subsequent studies in both cell signaling and cardiac electrophysiology models have demonstrated success in focusing parameter calibrations on optimizing prediction accuracy in place of specific parameter accuracy [11,48,49]. These analyses demonstrate the importance of calibrating models to not only fit available experimental data, but also generate accurate predictions of responses to novel perturbations.

Therefore, while optimizing the computational pipeline, we included a prediction metric in addition to parameter value accuracy and consistency. We created a quantifiable, pharmacologically-relevant measure of cell-specific arrhythmia susceptibility by finding the lowest level of I_Kr_ block that induced arrhythmic dynamics in each Kernik model (*in silico*) or iPSC-CM preparation (*in vitro*), which we termed the “I_Kr_ block tolerance threshold”. Calibrated models from our optimized computational pipeline were able to predict this threshold consistently and accurately, with similar error as those from calibrating to the original, non-normalized data.

The models calibrated to normalized data were also able to correctly rank the I_Kr_ block arrhythmia susceptibilities of the fitted Kernik model cells, suggesting that the models generated by this pipeline contain enough information to predict cell-specific arrhythmia susceptibilities, despite errors and non-constraint of several smaller conductance parameters.

### Limitations and potential directions

The model calibration process developed and optimized in this study can be applied broadly to research that investigates the ionic mechanisms behind iPSC-CM phenotype variability or predicts differential effects of therapeutics on various iPSC-CM cell lines and preparations. However, phenotypic variability also exists within iPSC-CMs from the same preparation. While it is possible to use fluorescent dyes to record electrophysiology of individual cells, we currently focused our efforts on calibrating models to represent multicellular preparations.

Future work could include adapting this computational pipeline to calibrate models representing single cardiomyocytes within an iPSC-CM preparation. Several conductance parameters fitted in our current pipeline did not contribute significantly to electrophysiological phenotype and responses (e.g. G_f_, G_SRleak_, and G_rel_), according to our parameter sensitivity analyses. Many of these same parameters were also less accurate and less constrained by our computational pipeline. While we showed that the resulting calibrated models could accurately predict several major electrophysiology phenotypes and channel block responses, we do not know whether these models could still predict responses that are strongly affected by one or more of the unconstrained parameters. Calibrated models generated by our computational pipeline should be evaluated on their ability to predict cell-specific susceptibility to other pro- or anti-arrhythmia triggers of interest, such as I_CaL_ or buffer [K^+^] changes. If improving parameter calibration accuracy and constraint of particular ionic current(s) is found to be necessary for model predictive power, one option would be to replace the conductances with lowest parameter sensitivity regression coefficient magnitudes with representative kinetic parameters, such as time constants or activation gates. If these kinetic parameters are found to impact electrophysiology in a manner that is detectable by the genetic algorithm, their calibrations will likely be more accurate and constrained, generating more precise cell preparation-specific iPSC-CM models.

## Conclusions

The optimized model calibration computational pipeline we developed in this study can be applied to ongoing pharmacological and disease studies to understand cell response variability in iPSC-CMs and advance precision medicine. Our pipeline simultaneously fits multiple model parameters to fluorescence voltage and calcium recordings, which are quicker and simpler to acquire than the patch clamp or other more involved electrophysiological techniques used in prior model calibration work. The ability to use fluorescence recordings increases this pipeline’s accessibility and throughput compared to prior methods. This unique characteristic makes our computational pipeline suitable for rapid generation of cell-specific models for therapeutic, disease, or population studies. Comparing cell-specific model parameters between iPSC-CM cell lines or to a benchmark, such as an adult cardiac model, would reveal ionic mechanisms behind this variability in iPSC-CM maturation and phenotypes. In addition, the personalized models created by this computational pipeline can simulate patient-specific drug and perturbation responses to quickly predict therapeutic efficacy or drug cardiotoxicity. In future studies, these calibrated iPSC-CM models can inform translation of ionic properties and physiological responses into adult cardiac models, which could, in turn, generate even more accurate, clinically-relevant cardiac electrophysiology phenotype and drug response predictions.

## METHODS

### Simulations of iPSC-CM electrophysiology

*In silico* simulations of iPSC-CM electrophysiology (e.g. membrane potential, calcium transients, changes in maximal conductances and ionic currents) were carried out in MATLAB R2020b by solving the ordinary differential equations defined in the Kernik et al. (2019) mathematical model of human iPSC-CMs [9], using the initial conditions listed in Supplementary Table S1. In cases where a protocol included cardiomyocyte pacing, electrical stimuli of 60 pA/pF were simulated for a duration of 1 ms at the specified intervals. All simulations were run for 5 minutes, which was sufficient to achieve a steady state using the baseline model and physiological, unperturbed conditions (Supplementary Table S3, protocol #19). The voltage and intracellular calcium concentration from the final 5 seconds of each simulation were stored and included in the *in silico* dataset.

### *In silico* dataset for optimization of iPSC-CM experimental protocol

To simulate a heterogeneous set of iPSC-CM cell lines, a “population” of Kernik model cells was created by drawing multiplier factors for each maximal conductance parameter from a log-normal distribution with mean multiplier = 1, spread = 0.2. This population was then simulated under physiological, unperturbed conditions (Supplementary Table S3, protocol 19), and filtered for cells which showed a spontaneous beating frequency between 0.3 Hz and 1.0 Hz, nearly matching the automaticity rate range for human iPSC-CMs reported in [50]. Out of the remaining model cells, 4 were randomly selected for simulation under the 19 different conditions, to generate the final *in silico* dataset. The Kernik model’s baseline conductance parameter values and the scaled conductances for these 4 model cells are listed in Supplementary Table S2. A list of the various conditions these 4 cells were simulated under can be found in Supplementary Table S3. All simulations were run for a 5-minute time span (steady state under physiological buffer conditions, without pacing), and membrane potential and intracellular calcium recordings from the final 5 seconds of these simulations were extracted and stored for the *in silico* dataset. The built-in *interp1* function in MATLAB was used to interpolate these data in equal time steps (0.1 ms). To mimic processed data from fluorescence recordings, we also created “normalized” versions of each dataset by scaling the data from a minimum of 0 to a maximum of 1. This dataset allowed for comparison of conductance parameter estimates produced during model calibration against known ground truth conductance values.

### Parameter calibration using the genetic algorithm

The data used for calibrating model parameters included the steady state AP and CaT waves from the previously-generated *in silico* dataset. Individual simulation runs are either included or excluded from the fitting procedure to compare different experimental “candidate protocols”. Rapid delayed rectifier potassium current (I_Kr_) block conditions were left out from protocol optimization to use for conductance estimate validation. If the candidate protocol consists of more than 1 condition, the final values of membrane potential, intracellular calcium concentration, and other steady state parameters are used as the starting values for the subsequent protocol condition. To determine the optimal protocol for accurate and consistent parameter identification, Kernik model maximal conductance parameters were fitted to *in silico* from each candidate protocol simulated for 4 of the dataset’s model cells, using the genetic algorithm (GA) [10,12]. Briefly, the GA creates a new population of Kernik model cells, each randomly assigned a set of conductance parameter scale factors, and simulates the candidate protocol for each of these model cells. Then, the mean squared error between corresponding data points in the GA-generated trace and the trace from the *in silico* dataset is calculated. The model cells with the lowest errors are retained, while higher-scoring cells have their parameter values altered in the next algorithm iteration. Details about specific GA settings, such as population size and parameter retention/alteration criteria, can be found in Supplementary Table S4. This process repeats for 20 iterations, where each new population has a lower average error than the previous. The parameters of the model cell with the lowest error from the final population are selected as the final calibrated parameter values from that run.

Parameter calibrations are conducted 10 times per candidate protocol per model cell, with each of the 10 GA runs starting with a different set of initial model populations. The 10 sets of conductance estimates from each GA run are evaluated on 1) accuracy to the ground truth parameter values of the model cells and 2) ability to predict the model cell’s response to a perturbation that was not seen during parameter calibration, such as I_Kr_ block.

### *In vitro* iPSC-CM tissue culture and optical recordings

Human induced pluripotent stem cells (iPSCs) derived from a single cell line were differentiated into cardiomyocytes (iPSC-CMs) by modulating canonical Wnt signaling [51]. The cardiomyocytes were enriched, then combined with cardiac fibroblasts in a ratio of 90% iPSC-CMs to 10% fibroblasts within a collagen-fibrin hydrogel solution (InvivoSciences, Inc., Madison, WI, USA). The engineered heart tissues (EHTs) thus formed were maintained in a serum-free cardiac maintenance medium, supplemented with penicillin and streptomycin (Thermo Fisher Scientific, Waltham, MA, USA), within 96-well micro culture plates (MC-96, InvivoSciences). EHT maintenance was carried out as previously described in relevant literature [52]. Following a five-day remodeling phase, the EHTs underwent further maturation with biphasic constant current electrical stimulations at a frequency of 1Hz for 8 days.

For optical voltage and calcium transient recordings, the EHTs were loaded with Fluovolt (at a dilution of 1:500, Thermo Fisher Scientific), or Cal-520 AM (also at 1:500, AAT Bioquest), along with PowerLoad (Thermo Fisher Scientific), by incubating them for one hour in Tyrode’s solution. This solution was prepared with either 1.0 mM or 1.8 mM [Ca^2+^]. Following the removal of the dyes, the solution was pH adjusted to 7.4 and warmed, and Tyrode’s solution with the corresponding calcium concentration was reintroduced to aid in recovery from dye loading stress over a 30-minute period. The EHTs were then paced at frequencies of 1.0 Hz, 1.25 Hz, or 2.0 Hz for five minutes. The steady-state membrane potential and intracellular calcium transients during this period were recorded using a high-throughput fluorescence plate imager, FDSS/µCell (Hamamatsu Photonics K.K., Japan), utilizing 470nm excitation and 540nm emission at a rate of 125 data points per second. The collected data were analyzed with the iVSurfer™ software (InvivoSciences), specifically designed for high-throughput waveform data analysis.

### Processing of *in vitro* iPSC-CM recordings for the computational pipeline

Fluorescence voltage and calcium recordings were processed in MATLAB with baseline drift subtraction (imerode function), median filtering to remove noise (medfilt1 function), and the same normalization steps as the pseudo-data. Kernik model maximal conductance parameters were fitted to these data using the same workflow as prior fittings to pseudo-data, with the 1.0mM buffer [Ca^2+^], 2.0Hz pacing data left out for validation.

## ABBREVIATIONS

AP: Action potential

CaT: Calcium transient

GA: Genetic algorithm

G_x_, I_x_, J_x_: Maximal conductance (G), current density (I), or flux (J) for x, where x can be: Na (Na^+^), f (funny Na^+^), CaL (L-type Ca^2+^), to (transient outward K^+^), Ks (slow delayed rectifier K^+^), Kr (rapid delayed rectifier K^+^), K1 (inward rectifier K^+^), PMCA (plasma membrane Ca^2+^ ATPase), bNa (background Na^+^), bCa (background Ca^2+^), Up (SR Ca^2+^ uptake), rel (SR Ca^2+^ release), NCX (Na^+^/Ca^2+^ exchanger), NaK (Na^+^/K^+^ ATPase), SRleak (SR leak), or CaT (T-type Ca^2+^)

iPSC-CM: Induced pluripotent stem cell-derived cardiomyocyte

LTCC: L-type calcium channel

NCX: Sodium-calcium exchanger

SERCA: Sarco/endoplasmic reticulum calcium ATPase

SR: Sarcoplasmic reticulum

